# AGAP1-associated endolysosomal trafficking abnormalities link gene-environment interactions in a neurodevelopmental disorder

**DOI:** 10.1101/2023.01.31.526497

**Authors:** Sara A. Lewis, Somayeh Bakhtiari, Jacob Forstrom, Allan Bayat, Frédéric Bilan, Gwenaël Le Guyader, Ebba Alkhunaizi, Hilary Vernon, Sergio R. Padilla-Lopez, Michael C. Kruer

**Author notes:** Sara A. Lewis, Somayeh Bakhtiari, Jacob Forstrom, Allan Bayat, Frédéric Bilan, Gwenaël Le Guyader, Ebba Alkhunaiz, Hilary Vernon, Sergio R. Padilla-Lopez. corresponding author: Michael C. Kruer.

## Abstract

*AGAP1* is an Arf1 GAP that regulates endolysosomal trafficking. Damaging variants have been linked to cerebral palsy and autism. We report 3 new individuals with microdeletion variants in *AGAP1*. Affected individuals have intellectual disability (3/3), autism (3/3), dystonia with axial hypotonia (1/3), abnormalities of brain maturation (1/3), growth impairment (2/3) and facial dysmorphism (2/3). We investigated mechanisms potentially underlying *AGAP1* neurodevelopmental impairments using the *Drosophila* ortholog, *CenG1a*. We discovered reduced axon terminal size, increased neuronal endosome abundance, and elevated autophagy at baseline. Given potential incomplete penetrance, we assessed gene-environment interactions. We found basal elevation in phosphorylation of the integrated stress-response protein eIF2α and inability to further increase eIF2α-P with subsequent cytotoxic stressors. *CenG1a*-mutant flies have increased lethality from exposure to environmental insults. We propose a model wherein disruption of *AGAP1* function impairs endolysosomal trafficking, chronically activating the integrated stress response, and leaving AGAP1-deficient cells susceptible to a variety of second hit cytotoxic stressors. This model may have broader applicability beyond *AGAP1* in instances where both genetic and environmental insults co-occur in individuals with neurodevelopmental disorders.

**Summary statement:** We describe 3 additional patients with heterozygous AGAP1 deletion variants and use a loss of function *Drosophila* model to identify defects in synaptic morphology with increased endosomal sequestration, chronic autophagy induction, basal activation of eIF2α-P, and sensitivity to environmental stressors.

## Introduction

Known genetic contributions to neurodevelopmental disorders are rapidly expanding and include both Mendelian and complex (non-Mendelian) phenomena. Mendelian inheritance patterns include classic autosomal dominant, recessive, and sex chromosome-linked monogenic disorders, currently estimated to lead to ~20% of autism spectrum disorders (Mahjani et al., 2021) and ~25% of cerebral palsy cases (van Eyk et al., 2021, Moreno-De-Luca et al., 2021). Autosomal dominant inheritance patterns represent important contributors to neurodevelopmental disorders, often arising spontaneously as *de novo* variants (Brunet et al., 2021) and may exhibit complex, non-Mendelian characteristics such as incomplete penetrance and variable expressivity (Ahluwalia et al., 2009). Other complex inheritance elements include epigenetic contributions to disease, gene-gene (epistatic) interactions and gene-environment interactions. Despite the inherent challenges in studying complex inheritance, such patterns represent important contributors to neurodevelopmental disorders including cerebral palsy (Fahey et al., 2017). Although substantial advances have been made in statistical and causal modeling of complex genetic factors (Madsen et al., 2011), few experimentally-validated genes and mechanisms have been described for complex inheritance leading to neurodevelopmental disorders such as cerebral palsy.

Missense variants in the Arf1-regulator *AGAP1* and heterozygous deletions in the 2q37.2 region that span this gene (Leroy et al., 2013) are associated with autism spectrum disorder (Cukier et al., 2014) and cerebral palsy (Chopra et al., 2022, Jin et al., 2020, van Eyk CL, 2019). Despite a statistical enrichment in *AGAP1*-variants in a cerebral palsy cohort (van Eyk et al., 2019), there is also evidence for environmental factors contributing to disease in several cerebral palsy patients with *AGAP1* variants. Therefore, the pathogenicity of these variants has not been definitively established and the disease-gene link for *AGAP1* is still unknown. Additionally, although the frequency is very low, not all individuals harboring *AGAP1* variants manifest neurodevelopmental disorders (gnomAD heterozygous LoF = 0.008%). Such incomplete penetrance is particularly common in individuals with autism spectrum disorder (Geschwind, 2011), and many autism susceptibility genes have thus been referred to as “risk genes” (Yuen et al., 2017). To better understand potential contributors to the apparent complex disease relationship of *AGAP1*, we sought to identify underlying pathophysiology and characterize potential gene-environment interactions caused by *AGAP1* loss of function.

AGAP1 is broadly expressed with RNA levels highest in brain and protein expression highest in lung, endocrine tissues, male reproductive tissue, adipose, and bone marrow/lymphoid tissues (https://www.proteinatlas.org/ENSG00000157985-AGAP1/tissue). AGAP1 is expressed in the mouse and human brain throughout pre- and post-natal development, with subcellular localization to dendrites, axons, and synapses (Arnold et al., 2016). This suggests AGAP1 plays a role in brain development, although studies of its function are limited. *AGAP1* is predicted to be intolerant to LoF variants (observed/expected LoF = 0.1169; LoF Z score = 5.864; pLI = 1.0) and potentially missense variants as well (observed/expected missense ratio = 0.8479; missense Z score = 1.287). AGAP1 contains a GTPase like domain (GLD), PH domain, and GAP-activity domain (http://www.ebi.ac.uk/interpro/protein/UniProt/Q9UPQ3/). The GAP activity of AGAP1 activates Arf1 hydrolysis to complete the GTPase cycle (Nie et al., 2003). The proper initiation and termination of Arf1 signaling is required for actin cytoskeleton polymerization (Saila et al., 2020, Davidson et al., 2015) which drives the formation of vesicles (Heuvingh et al., 2007). Overexpression of AGAP1 creates punctate endocytic structures and reduces stress fibers, demonstrating potential roles in regulating actin dynamics and trafficking from the endocytic compartment (Nie et al., 2003).

AGAP1 also regulates protein trafficking via the PH domain by recruiting subunits σ3 and δ of the AP-3 protein trafficking adaptor complex. AP-3 positively regulates the movement of proteins to the lysosome (Nie et al., 2003) and plasma membrane (Bendor et al., 2010). Immunostaining confirms AGAP1 localization to early and recycling endosomes (Arnold et al., 2016) and *Golgi* (Cukierman et al., 1995). Both increases and decreases in AGAP1 levels decrease the rate of protein trafficking out of Rab11-positive endosomes (Arnold et al., 2016) and AGAP1-mediated endocytic recycling regulates M5 muscarinic receptor surface localization and consequently, neuronal function (Bendor et al., 2010). AP-3 is important for trafficking lysosome membrane proteins, such as LAMP1, from *Golgi* to lysosome (Chapuy et al., 2008). Therefore, AGAP1 is predicted to be a key regulator of protein trafficking from the early endosome to the *Golgi*, lysosome, or plasma membrane.

To better understand how AGAP1 may contribute to nervous system function and expand on the potential *AGAP1*-disease link, we report clinical phenotypes of 3 individuals not previously reported with microdeletion variants and compare phenotypes with prior reports. We have previously described impaired locomotion from a loss of function variant of *CenG1a*, the *Drosophila* ortholog to *AGAP1* (Jin et al., 2020). *CenG1a* has well-conserved protein domains (Gündner et al., 2014) (DIOPT 13/16) and *Drosophila* models have been widely used to dissect mechanisms of neurological disease (Ugur B, 2016). Here we investigated the role of *CenG1a* in neuronal morphology, endo-lysosomal distribution and composition, autophagy, and stress responses in this genetic model.

## Results

### Patient clinical phenotypes

DOMINO predicts that heterozygous variants in AGAP1 will lead to haploinsufficiency (P(HI) 0.8548). AGAP1 is also intolerant to genomic loss-of-function variants (pLI 0.9994). These observations provide complementary evidence for their role in AGAP1-associated neurodevelopmental disorders.

We identified 3 new patients heterozygous for copy number variants that led to complete or partial AGAP1 gene deletion (exons 7-9 in one patient and exons 10-13 in another) (**Table 1**). We noted phenotype overlap with 6 previously reported patients with AGAP1 variants or 2q37 deletions limited to the AGAP1 region (**Table 2**). Key features include intellectual disability/developmental delay, language impairments, autism spectrum disorder, aggression, and impaired length, weight, and cranial growth (**Fig. 1**). With lower frequency, we also observed epilepsy, dystonia and/or axial hypotonia, brain growth abnormalities, eye abnormalities, and skeletal defects. The shared clinical features of this expanded cohort provide additional support that *AGAP1* variants can lead to a mixed neurodevelopmental disorder with systemic manifestations.

**Figure 1:**
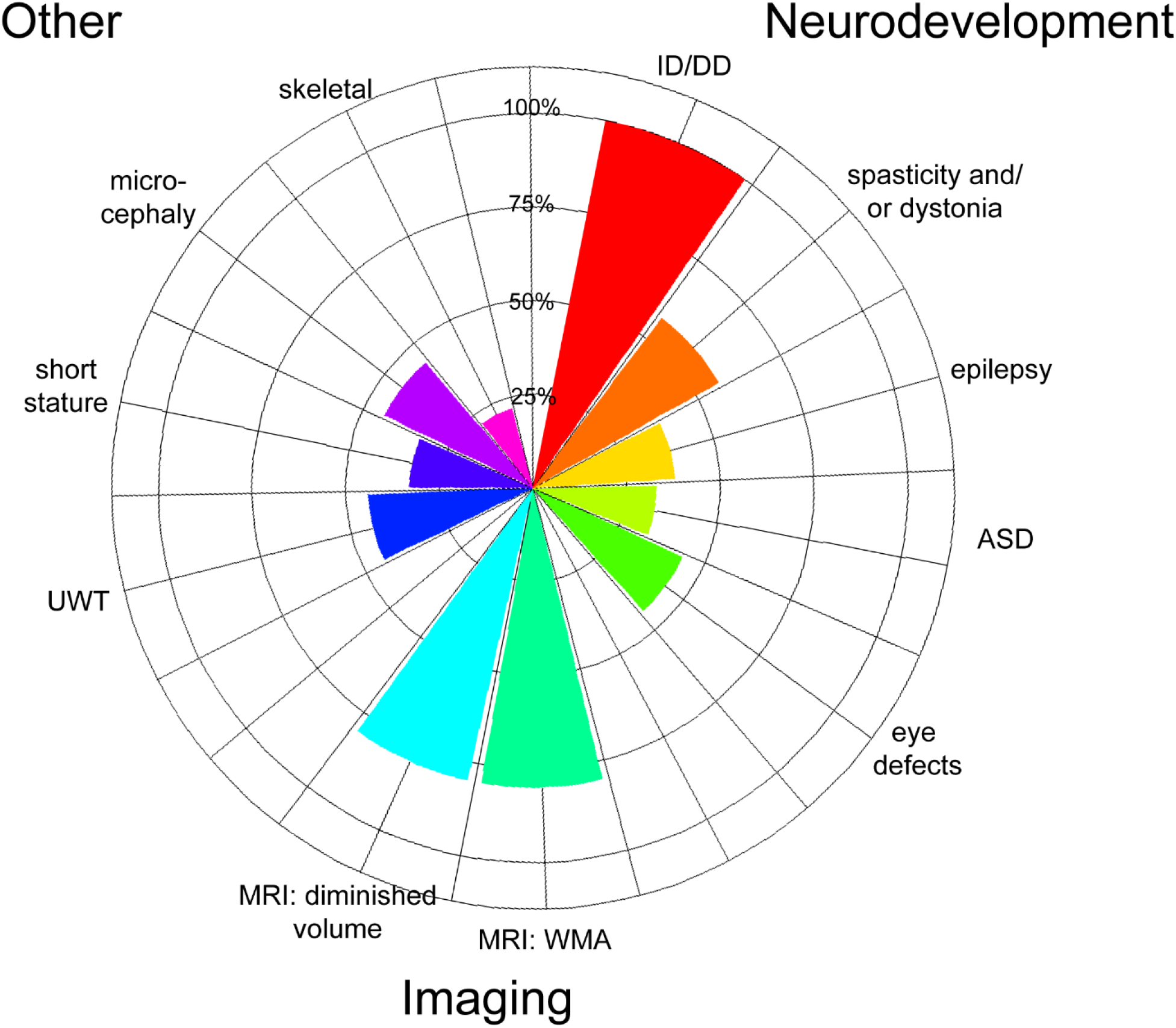
Phenogram of shared features from 10 AGAP1 patients. 3 patients from this study and 6 previously reported patients with heterozygous, deleterious, Exac=0 variants (van Eyk CL, 2019) (Jin et al., 2020) (Leroy et al., 2013) (Pacault et al., 2018) (Chopra et al., 2022) WMA=white matter abnormalities. UWT=underweight. Denominators based on number of patients with phenotype reported in the publication. MRI findings calculated using 5 patients with MRI studies.

**Table 1:** Clinical phenotypes of newly reported AGAP1 patients. Domains assessed using Uniprot: GTPase domain p. 66-276 and PH (pleckstrin homology; AP3 binding) p.346-588 and Arf-GAP p. 609-729.

|  |  |  |  |
| --- | --- | --- | --- |
| **variant(s)** | 2q37.2 deletion *AGAP1* only | microdeletion exons 7 to 9 | microdeletion exons 10 to 13 |
| **affected domain** | whole gene | GTPase | PH |
| **CADD score** | - | - | - |
| **inheritance** | maternal | de novo | paternal |
| **age** | 4 y | 7 y | 13 y |
| **sex** | M | F | M |
| **DD/ID** | DD | some DD | profound ID |
| **autism/language defects** | ASD; delayed speech | language defects | language absent |
| **epilepsy** | none | none | none |
| **movement disorder** | none | none | psychomotor delay, axial hypotonia and peripheral hypertonia esp. in lower limbs |
| **eye disorders** | none | none | none |
| **behavioral disorder** | none | none | severe aggression |
| **skeletal disorders** | none | clubfoot, short long bones, hyperlaxity, pectus excavatum, | none |
| **craniofacial features** | high forehead, bi-temporal narrowing, depressed nasal bridge, anteverted | elongated face with a tubular nose, a narrow mouth, columella salient, | none |
|  | nostrils, ears pointy and protrude | ears have transverse fold, palmar transverse fold |  |
| **obesity or growth** | normal weight (90 centile) and length (39 centile) | height 37.5 cm (-1.7 SD) @ birth and (-1 SD) @ 7y. normal weight | height and weight <2SD |
| **early mortality** | none | none | none |
| **brain structure and/or microcephaly** | normal OFC (48.5 cm 10 centile @ 18 m) | microcephaly (OFC 27 cm -2 SD @ birth and -1.5 SD @ 7y) normal MRI | microcephaly (OFC <3SD) |
| **History of adverse event** | None | Yes (premature rupture of membranes with premature birth (31 weeks gestation), neonatal jaundice) | Not reported |

**Table 2:** Quantification of clinical phenotypes from patients with heterozygous, deleterious variants. Variants from this study and previously published organized by affected domain. Phenotypes overlapping with model organism findings: M=mouse (mousephenotype.org), z=zebrafish (van Eyk et al., 2019), f=fly (this study; Jin et al., 2020; Gunder et al., 2014). TTTS=twin-to-twin transfusion syndrome.

| Neurological | GTPase | PH | Both | % total |
| --- | --- | --- | --- | --- |
| DD/ID<sup>z,f</sup> | 1/1 | 4/4 | 4/4 | 9/9 (100%) |
| brain abnormalities and/or microcephaly | 0/1 | 4/4 | 2/5 | 6/9 (67%) |
| Eye disorders (nystagmus/extropia) | 0/1 | 2/4 | 2/4 | 4/9 (44%) |
| spasticity (diplegia or quadriplegia)<sup>z,f</sup> | 0/1 | 3/4 | 1/4 | 4/9 (44%) |
| Epilepsy | 0/1 | 2/4 | 1/4 | 3/9 (33%) |
| Autism | 1/1 | 0/4 | 2/4 | 3/9 (33%) |
| dystonia w/ or w/out axial hypotonia<sup>z,f</sup> | 0/1 | 1/4 | 1/4 | 2/9 (22%) |
| behavioral disorder | 0/1 | 1/4 | 1/4 | 2/9 (22%) |
| Other |  |  |  |  |
| short stature/under-weight/failure to thrive | 0/1 | 2/4 | 2/4 | 4/9 (44%) |
| skeletal disorder<sup>m</sup> | 1/1 | 0/4 | 2/4 | 3/9 (33%) |
| craniofacial features<sup>m</sup> | 1/1 | 0/4 | 1/4 | 2/9 (22%) |
| early mortality<sup>m</sup> | 0/1 | 0/4 | 1/4 | 1/9 (11%) |
| obesity | 0/1 | 0/4 | 1/4 | 1/9 (11%) |
| Cardiac abnormalities | 0/1 | 0/4 | 0/4 | 0/9 (0%) |
| Variant | Domain | Inheritance | Patient History | Reference |
| 1 c.1400C>G p.P467R | PH | De novo | born 28 weeks,<br>TTTS | van Eyk et al. 2019 |
| 2. c. 1232C>T p.P411L | PH | Not maternally inherited | Prematurity, intraventricular haemorrhage | van Eyk et al. 2019 |
| 3 c.1462G>A p.Asp488Asn | PH | Not maternally inherited | non-cryptogenic | Chopra et al. 2022 |
| 4 c.957+1G>A splicing | both | De novo | Emergency caesarean with APGAR scores 1@ 1 min, 7 @ 5 min | Jin et al., 2020 |
| 5 deleted region<br>Chr2:235670886–<br>236827204 | both | Not available | Not available | Leroy et al., 2013 |
| 6 deleted region<br>chr2:234966458-236025317 | both | De novo | born full term after an uneventful pregnancy | Pacault et al., 2019 |

### CenG1a is required for scaled growth of the neuromuscular junction

We noted movement disorders such as dystonia in *AGAP1* patients (**Table 2**) and previously showed locomotor impairments in *CenG1a* mutant flies. *AGAP1* patients also exhibit microcephaly, impaired cortical growth and delayed myelination and *AGAP1* is known to be important for dendritic spine maturation (Arnold et al., 2016). We therefore screened CenG1a mutants for neuroanatomical phenotypes. We utilized a null allele (*CenG1a^Δ9^*) given that AGAP1 putatively acts via loss of function. We found a reduction in the size of the axon terminal (**Fig. 2B**) without a corresponding decrease in the muscle area in *CenG1a^Δ9^* homozygotes; the *CenG1a^Δ9^/Df* hemizygotes had an unexpected increase in muscle size (**Fig. 2C**). We did not detect a change in bouton number or density (number/neuromuscular junction (NMJ) area) or number of satellite boutons (**Fig. 2D**). Other than the decreased size, the neuron appeared morphologically normal with no change in branch number (**Fig. 2E**). Pre- and post-synaptic marker colocalization was normal (not shown). A degenerative phenotype often has post-synaptic signal with absent presynaptic signal (Lincoln et al., 2015). In contrast, a synaptic maturation phenotype often has presynaptic signal with absent postsynaptic signal (Vasin et al., 2014). Given that *CenG1a* mutants demonstrated reduced synapse size, our findings indicate this likely represents an impaired neuronal growth or remodeling phenotype rather than representing degeneration or defective synaptogenesis.

**Figure 2:**
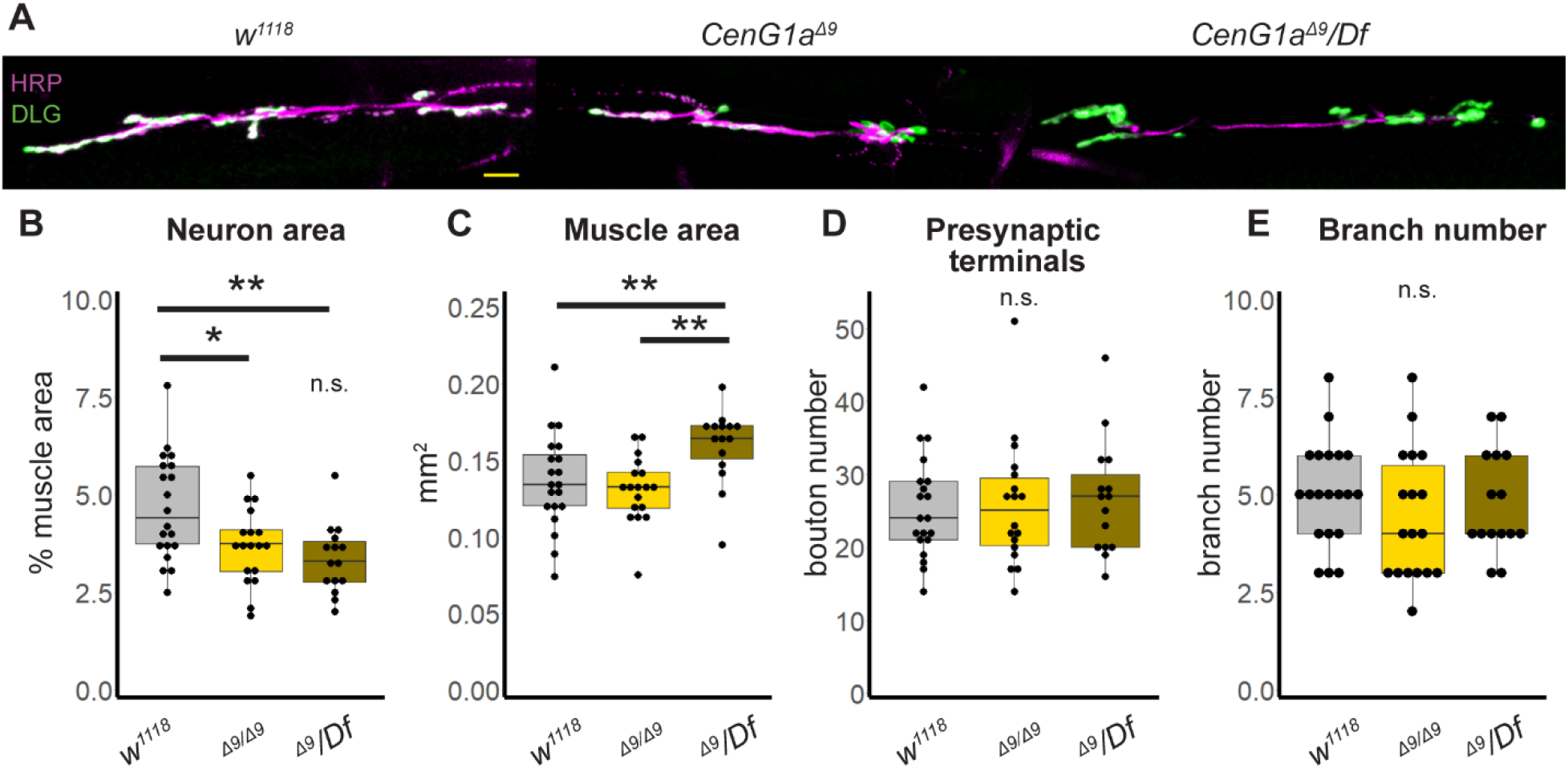
AGAP1 regulates scaled growth at the Drosophila NMJ. A. Images of axon terminals from late 3^rd^ instar larva. HRP (magenta) marks neuron membrane and DLG (green) marks postsynaptic structures to identify individual boutons. Scalebar=20 μm. B-E. Box and whisker plots of genetic control (*w^1118^*), *Ceng1a* null homozygote (*Δ9/Δ9*) and *CenG1a* hemizygote (*Δ9/Df*) to map phenotype to CenG1a genetic locus. Neuron (HRP) area normalized to muscle area is significantly decreased in both *CenG1a* mutants (B). Muscle area is also increased in hemizygotes, but not in homozygotes (C). The number of boutons, i.e. presynaptic terminals, (D) and axon branches (E) are not changed in *CenG1a* mutants. *Df=Df(2L)BSC252* is a deletion uncovering *CenG1a*, NMJ=neuromuscular junction, HRP= horseradish peroxidase (neuron membrane), DLG=discs large, psd95 ortholog Boxes represent 25^th^ and 75^th^ percentiles; whiskers represent 10^th^ and 90^th^ percentiles. *w^1118^* n=20 NMJ, 13 animals. *CenG1a^Δ9^* n=18 NMJ, 10 animals. *CenG1a^Δ9^/Df* n=15 NMJ, 10 animals. * p<0.05, ** p<0.005 by 2-tailed Mann-Whitney rank sum test.

### Increase in number and lysosomal localization of Rab7-positive endosomes in motor neurons of CenG1a mutants

AGAP1 is predicted to regulate protein trafficking out of the endosome (Arnold et al., 2016, Nie et al., 2003). We investigated whether *AGAP1* loss of function disrupted trafficking within the endosomal compartment in the nervous system using the *CenG1a* knockout model. We examined the Rab7 marker of early through late endosomes (Shearer and Petersen, 2019) in the NMJ (**Fig. 3A-B**), using HRP-defined area to distinguish between axon terminals and muscles. We noted increased Rab7 signal in *CenG1a* mutants (**Fig 3B’**) compared to genetic controls (**Fig 3A’**), particularly within HRP-defined neuronal area. The area (**Fig. 3C**) and density of Rab7-positive puncta (**Fig. 3C’’**) was increased in *CenG1a* mutants with no change in average size (**Fig. 3C’**) intensity of signal (not shown). The increased endosome signal was localized to the motor neuron with no change in muscle, consistent with nervous system-specific expression of *CenG1a*. Thus, *AGAP1/CenG1a* appears to be important for regulating the number and size of endosomes in neurons.

**Figure 3:**
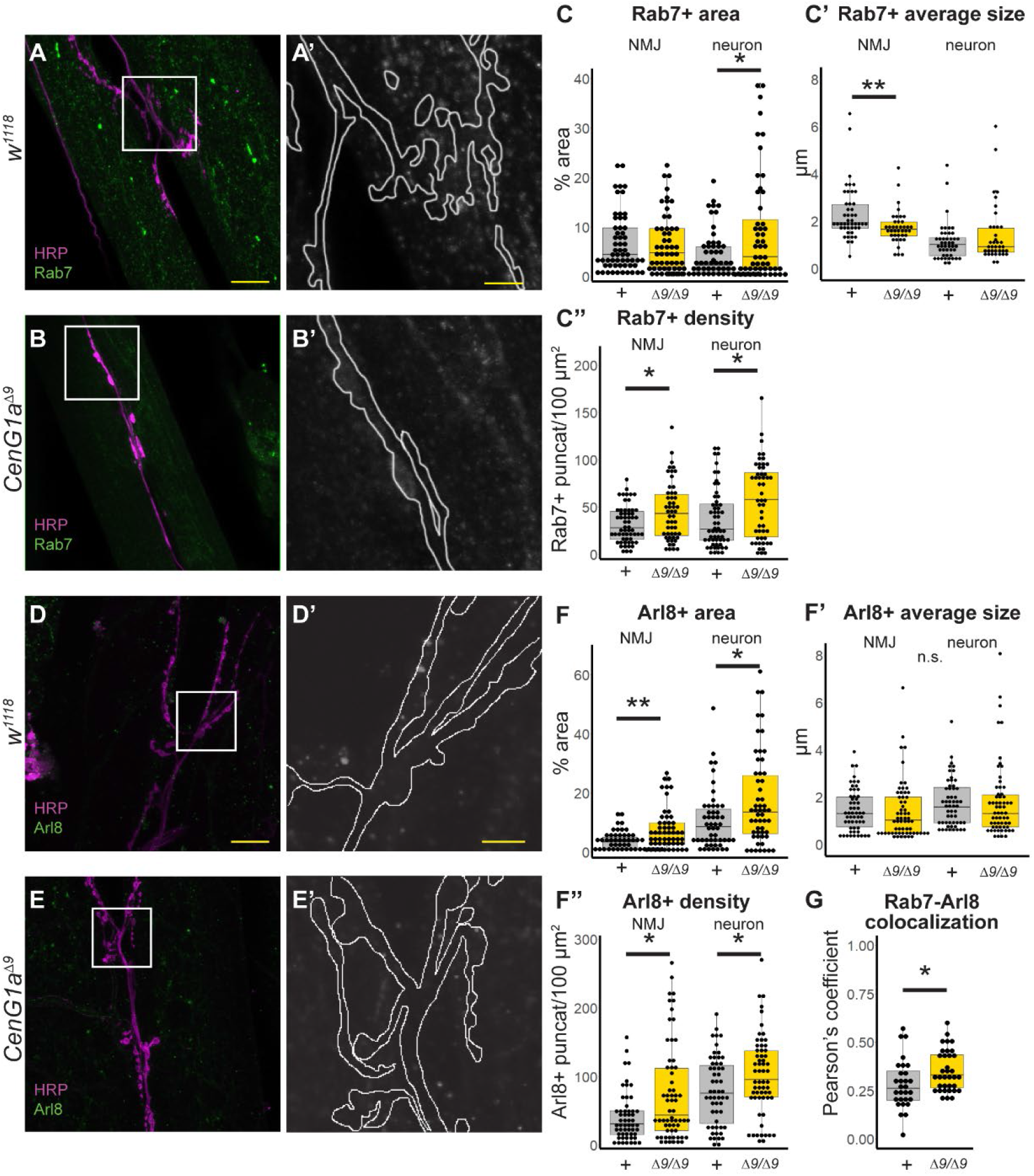
AGAP1 regulates the endosomal and lysosomal compartments at the Drosophila NMJ. A-B. HRP (magenta) localizes to neuron membranes and Rab7 (green) is present in early to late endosomes of *w^1118^* controls (A) and *CenG1a^Δ9^* null homozygotes (B). Scalebar=20 μm. A’-B’. Magnified view of box from panels A and B showing the Rab7 channel with the neuron outlined. Notably more Rab7 staining is present in the outlined neuronal area in CenG1a-mutant NMJs (B’) compared to controls (A’). Scalebar=5μm. C. Quantification of Rab7 puncta properties. The area (C’), and number of puncta per area density (C’’) are increased in the neurons of *CenG1a^Δ9^* loss-of-function mutants, but not endosome-associated average puncta size (C’’). +=*w^1118^*, Δ9=*CenG1a^Δ9^*. D-E. HRP (magenta) and Arl8 (green), a marker of lysosomes in controls (D) and *CenG1a^Δ9^* null homozygotes (E). Scalebar=20 μm. D’-E’. Magnified view of box from panels D and E showing the Arl8 channel with the neuron outlined. Notably more Arl8 staining is present in the outlined neuronal area of CenG1a-mutant NMJs (E’) compared to controls (D’). Scalebar=5 μm. F. Quantification of Arl8 properties. The lysosome-associated area (F) and density (F’’) is increased in neuron and, to a lesser extent, across the entire NMJ in *CenG1a^Δ9^* mutants while the average size is unaffected (F’). G. Pearson correlation coefficient is significantly increased between Rab7- and Arl8-positive puncta in *CenG1a^Δ9^* mutant neurons. NMJ=neuromuscular junction, HRP=horseradish peroxidase, fov=field of view. Numbers for 1C (Rab7): *w^1118^* n=58 fov, 20 NMJ, 12 animals. *CenG1a^Δ9^* n=52 fov, 21 NMJ, 10 animals. Numbers for 1F (Arl8): *w^1118^* n=51 fov, 23 NMJ, 14 animals. *CenG1a^Δ9^* n=54 fov, 25 NMJs, 14 animals. Numbers for 1F colocalization: *w^1118^* n=29 fov, 13 NMJ, 8 animals. *CenG1a^Δ9^* n=35 fov, 15 NMJ, 8 animals. *** p<0.0005, ** p<0.005, * p<0.05 by 2-tailed Mann-Whitney rank sum test.

We investigated whether increased endosome density was due to accumulated endosomes resulting from a failure of macroautophagy, reflecting disrupted endo-lysosomal degradation. We examined Arl8, a marker of lysosomes (Rosa-Ferreira et al., 2018) and noted it was also increased in *CenG1a*-mutant NMJs (**Fig. 3E’**) compared to genetic controls (**Fig. 3D’**). The lysosome-associated area (**Fig. 3F**) and density (**Fig. 3F’’**) of Arl8 was increased in *CenG1a* mutants, although the number of lysosomes was unchanged (**Fig. 3F’’**). We subsequently found that there was an increased colocalization between Rab7 and Arl8 puncta in the *CenG1a* mutants (**Fig. 3G**), reflecting an increased endo-lysosomal compartment. Altogether, this data demonstrates increases in average lysosome size and content, potentially from increased endosomal fusion (Jacomin et al., 2016), a defect in lysosomal degradation, or disrupted endo-lysosomal reformation (Klionsky et al., 2008).

**Figure 4:**
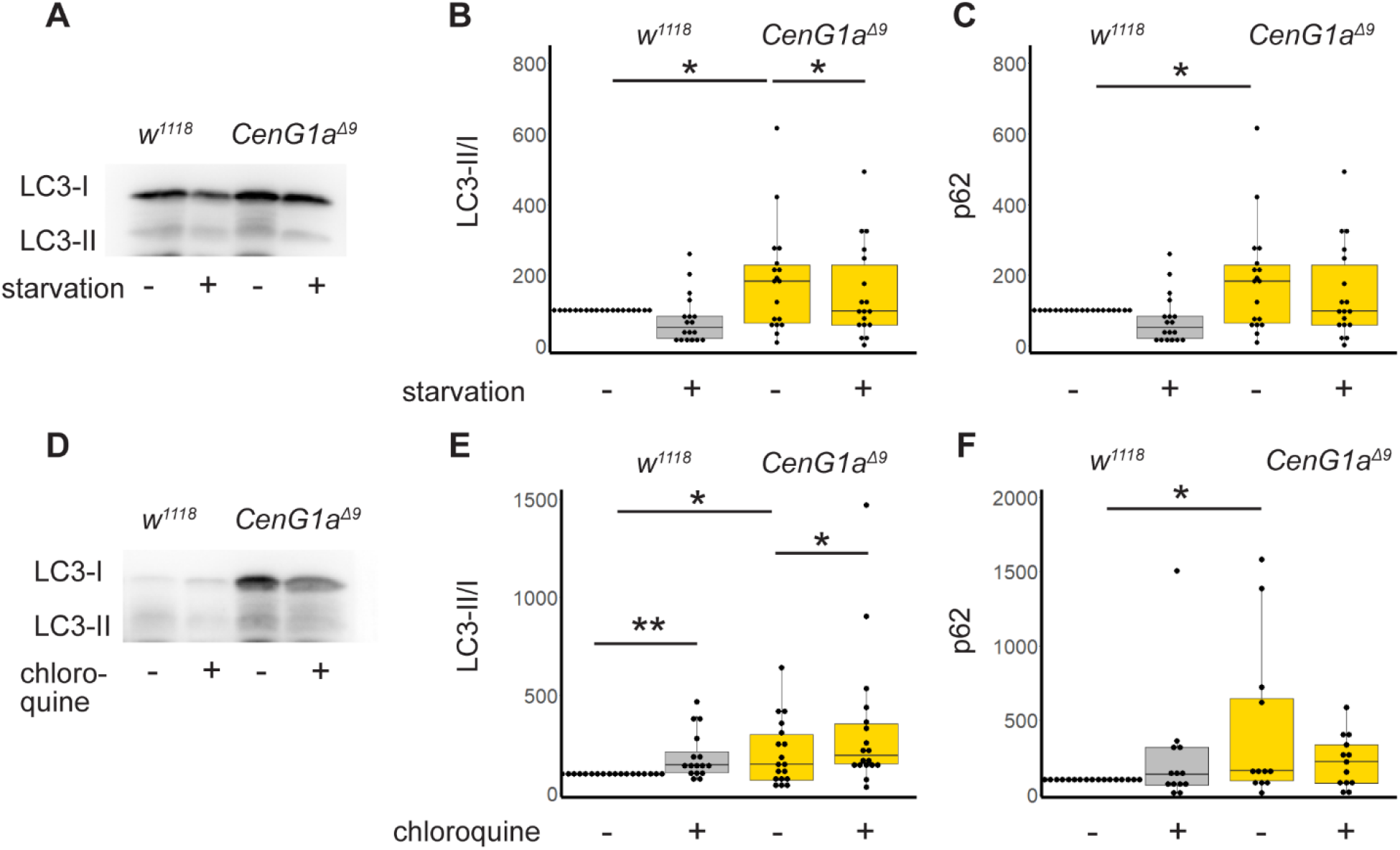
CenG1a mutants have increased autophagy induction with normal flux. A. Representative western blot of LC3-II and LC3-I (Atg8a) from *w^1118^* controls and *CenG1a^Δ9Δ9^* homozygous mutants from either fed or starved conditions. B. Quantification of LC3-II/I ratio from larva with or without starvation. Values normalized to within experiment *w^1118^* fed condition. Fed *CenG1a^Δ9^* animals have increased LC3-II ratio compared to fed controls. Control LC3-II ratios do not change after 24 hours of starvation while *CenG1a^Δ9^* mutants decrease to genetic control levels. n=17 from 6 biological replicates of 20+ larva. C. Ref2(P)/p62 from larva with or without starvation. Ref2(P) levels are elevated in *CenG1a^Δ9^* and do not change during starvation for either genotype. n=19 from 5 biological replicates. D. Representative blot of LC3-II and LC3-I (Atg8a) with 24 hours feeding with vehicle control or chloroquine. E. Quantification of LC3-II/I ratio from larva with or without chloroquine feeding to block autophagic flux. *CenG1a^Δ9^* animals have elevated LC3-II/I ratios compared to controls at baseline. Chloroquine feeding elevated LC3-II/I ratios in both genotypes, demonstrating normal autophagic flux in *CenG1a* mutants. n=10 from 5 biological replicates. F. Ref2(P) levels are elevated in *CenG1a^Δ9^* mutants at baseline, and levels do not change during 24 hours of chloroquine feeding. n=17 from 6 biological replicates. Values normalized to within experiment *w^1118^* vehicle control condition. * p<0.05 by 2-tailed paired t-test. All other comparisons n.s.

### CenG1a mutants have an increased rate of basal autophagy but normal flux

We next asked whether the increased endosomal-lysosomal colocalization was due to impaired autophagic flux or increased autophagy in *CenG1a^Δ9^* mutants. We examined Atg8a, the *Drosophila* ortholog of LC3, which is lipidated in order to target membranes to the autophagosome. This modification is detected as a size shift in western blotting and calculated as lipidated/basal (LC3-II/I) levels. We found *CenG1a* has a higher ratio of LC3-II/LC3-I compared to genetic controls at baseline (**Fig 4B**). We induced starvation to assess the ability of *CenG1a^Δ9^* mutants to respond to an exogenous cytotoxic stressor. LC3-II/LC3-I ratios did not change in genetic controls after 24 hours of starvation, indicating optimized balance between autophagy induction and protein degradation and clearance. In contrast, starvation decreased the basally elevated LC3-II/LC3-I ratios in *CenG1a* mutants.

We then tested whether autophagic flux was normal by treating larva with chloroquine (Zirin et al., 2013), a weak base that blocks autophagy by altering the acidic environment of lysosomes and autophagosome-lysosome binding. We found chloroquine feeding increased ratios of LC3-II/LC3-I for both genetic controls and *CenG1a^Δ9^* mutants, as would be expected when blocking normal autophagic flux (**Fig. 4E**). This also suggests that lysosomal function is unaffected in *CenG1a* mutants.

We then assessed Ref(2)P, the *Drosophila* ortholog of p62, which recruits ubiquitinated cargos to autophagosomes; p62 accumulates when autophagy is impaired (Pircs et al., 2012). Consistent with elevated LC3-II/LC-I ratios, we found increases in Ref(2)P in *CenG1a^Δ9^* larva compared to genetic controls at baseline (**Fig. 4C, 4F**). We did not detect a significant difference in Ref(2)P within genotypes between fed and starved conditions nor with chloroquine treatment at 24 hours. Together, our data demonstrate higher levels of autophagy induction in *CenG1a* mutants at baseline, but with preserved flux and increased clearance during starvation.

### CenG1a mutants have a diminished capacity to respond to additional stressors via eIF2α phosphorylation

Disruptions of protein trafficking (Amodio et al., 2019) and clearance (Xiong R, 2013) have been linked to activation of the integrated stress response. When the alpha subunit of the eukaryotic initiation factor-2 (eIF2α) is phosphorylated, global protein translation decreases while the expression of stress response genes and autophagy components is selectively upregulated (Humeau et al., 2020, B’chir et al., 2013). We therefore investigated whether loss of *CenG1a* altered eIF2α phosphorylation (eIF2α-P).

We found a significant increase in phosphorylated eIF2α *CenG1a^Δ9^* in larva and young, 1-day-old adults (**Fig. 5B**). Control adults increased phosphorylation between 1 and 14 days post-eclosion, consistent with previous descriptions of normal aging (Lenox et al., 2015). In contrast, *CenG1a^Δ9^* animals showed elevated phosphorylation early in development but mutant animals did not show an age-dependent upregulation of eIF2α-P over time. These results indicate that eIF2α phosphorylation is chronically activated in mutant animals.

**Figure 5:**
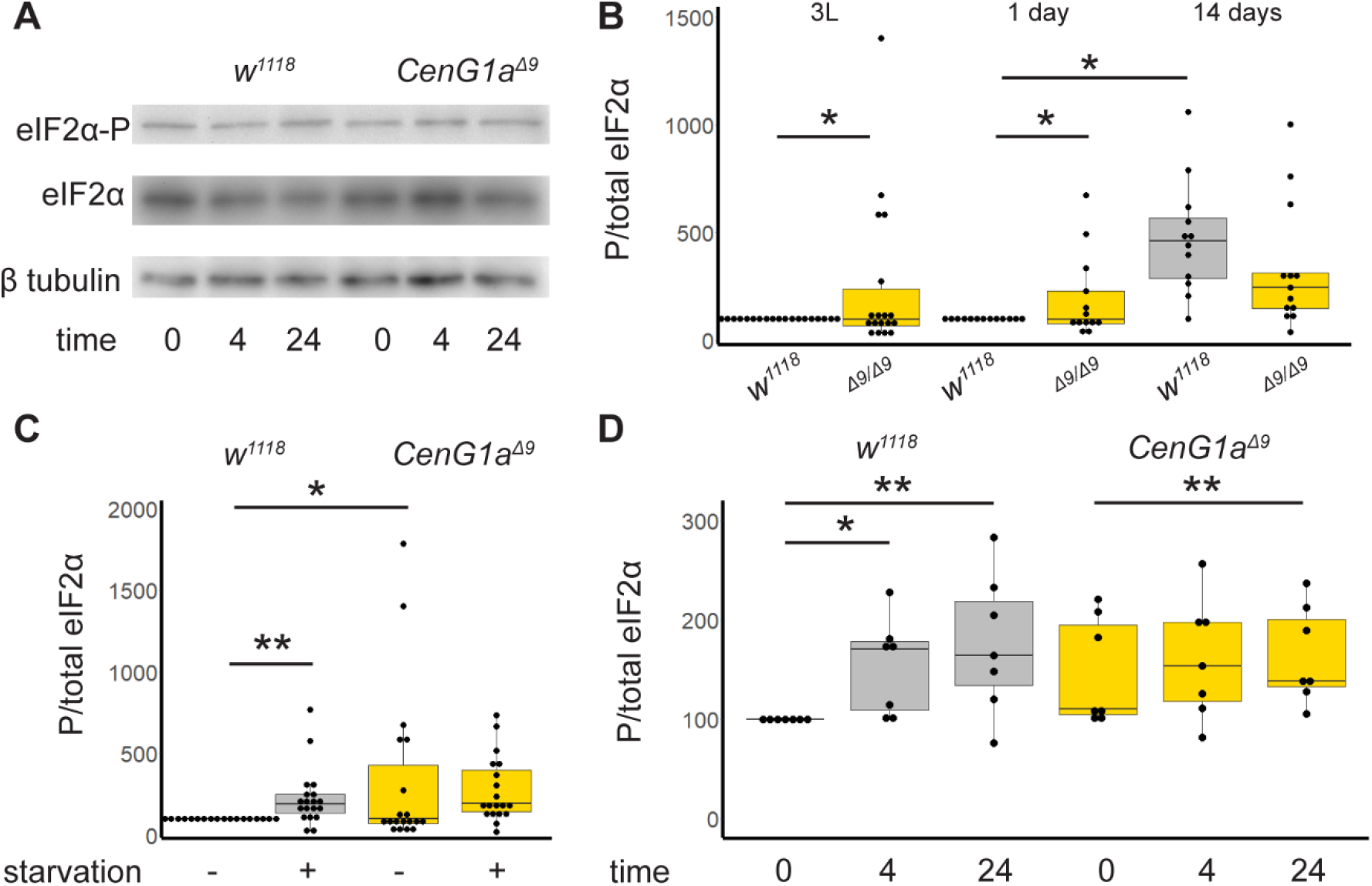
CenG1a mutants have elevated levels and defective regulation of eIF2α phosphorylation. A. Representative western blots from 3 day old adults exposed to tunicamyocin for 0, 4, or 24 hours, cropped to show phosphorylated eIF2α (eIF2α-P) and then the same blots reprobed for total eIF2α levels and β tubulin. Quantification shown in D.B. eIF2α-P of *w^1118^* (grey) and *CenG1a^Δ9^* homozygous (gold) larva (left), 1 d.o. (center) and 14 d.o. (right). Control animas have a singificant increase in eIF2α-P from 1 to 14 days post-eclosion. In contrast, *CenG1a* mutants do not alter eIF2α-P levels with age and have reduced eIF2α-P compared to wildtype controls at 14 days post eclosion. n=14 from 5 biological replicates for larva and n=13 from 3 biological replicates for adults. C. eIF2α-P of 2^nd^ instar larva either deprived of yeast (starved) or fed for 24 hours before protein extraction. Wildtype flies increase eIF2α-P in response to starvation stress while *CenG1a^Δ9^* mutants have elevated eIF2α-P that fails to change in response to starvation. n=14 from 5 biological replicates. D. Adults 3 days post eclosion were moved to vials with tunicamycin-containing food (12 μM) for 0, 4, or 24 hours before protein extraction. Control animals increase eIF2α-P in response to tunicamycin-induced unfolded protein stress at 4 and 24 hours. In contrast, *CenG1a^Δ9^* mutants are equivocally elevated at baseline (p=0.06) and have delayed response where eIF2α-P does not increase until 24 hours of exposure. n=7 from 2 biological replicates. Baseline data shown in B taken from C and D. Values normalized to within experiment *w^1118^* no treatment condition. * p<0.05 ** p<0.005 by 2-tailed paired t-test. All other comparisons n.s.

We next examined whether basal elevations in autophagy and eIF2α-P impaired responses to stress in a second hit scenario. Amino acid starvation normally activates phosphorylation of eIF2α through the GCN2 kinase (Pakos-Zebrucka et al., 2016). When challenged with 24 hours of starvation, control larva exhibited increased eIF2α phosphorylation. In contrast, *CenG1a^Δ9^* larva with chronically elevated eIF2α-P failed to increase it further (**Fig. 5C**). This indicates a diminished capacity to respond to a 2^nd^ hit cytotoxic stressor.

This suggested to us that *CenG1a^Δ9^* animals may have increased sensitivity to exogenous stressors. Accordingly, we exposed adult flies to tunicamycin which blocks protein glycosylation. Tunicamycin impairs protein folding, induces the integrated stress response in the ER, and triggers phosphorylation of eIF2α via PERK (Chow et al., 2013). Control animals increase eIF2α phosphorylation (eIF2α-P) at 4 and 24 hours of tunicamycin exposure. In contrast, eIF2α-P was equivocally upregulated in *CenG1a*-mutant adults compared to controls at baseline [p=0.06]. In this context, *CenG1a* mutants exhibit a delay in their activation of stress response pathways; there was a failure to increase eIF2α-P at 4 hours, although phosphorylation was increased after 24 hours (**Fig. 5D**). Thus, multiple stress-response pathways are chronically active in *CenG1a* mutants and do not respond appropriately to acute stressors.

We confirmed elevated eIF2α-P has functional consequences on the rates of protein synthesis with puromycin labeling. As expected, control larva decreased global protein synthesis in the presence of tunicamycin-induced ER stress. In contrast, *CenG1a^Δ9^* mutants have decreased levels of protein synthesis at baseline and fail to further reduce levels of protein synthesis in the presence of this stressor (**Fig. 6A,B**).

**Figure 6.**
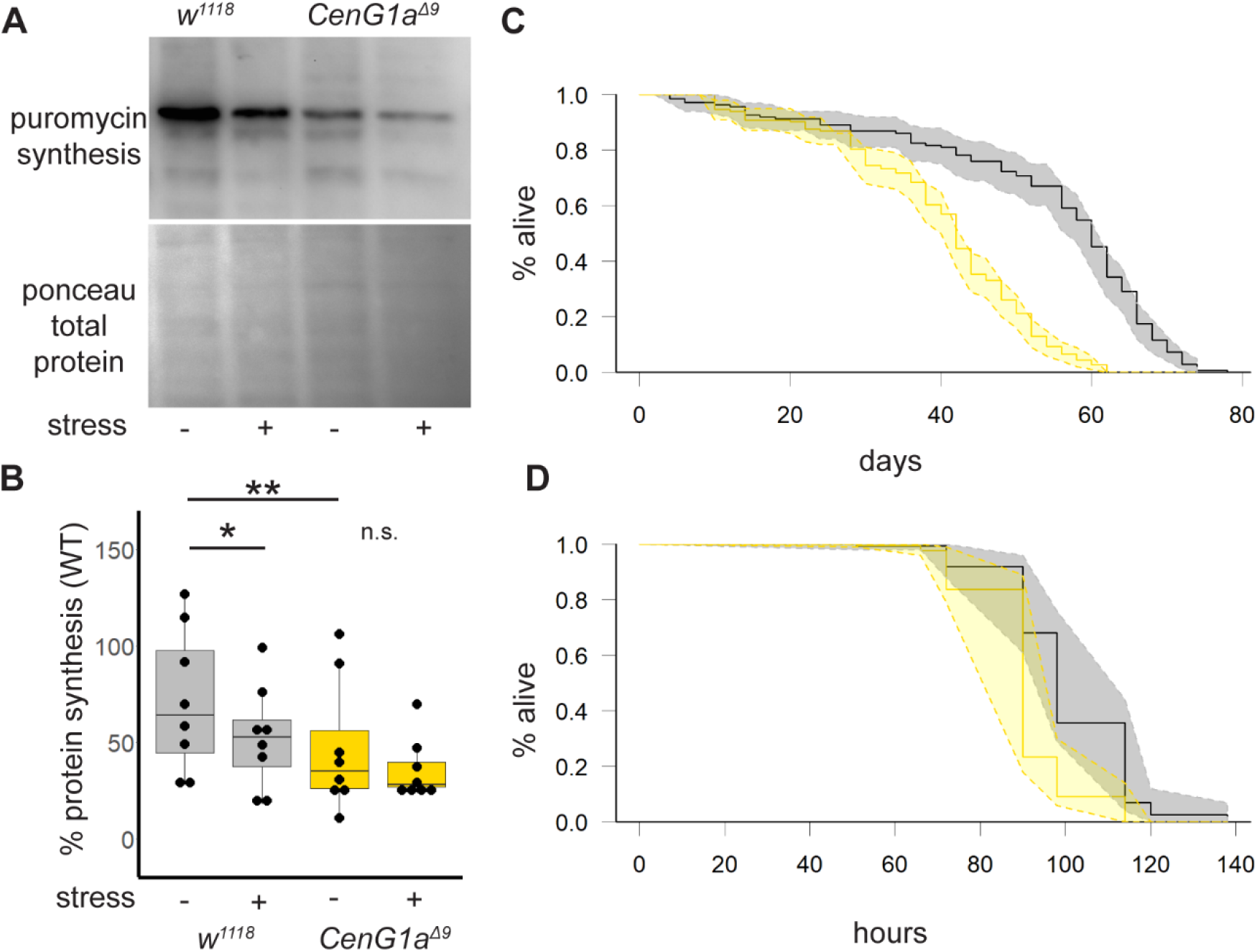
Decreased protein translation and sensitivity to stress in CenG1a mutant flies. A. Newly synthesized proteins identified by puromycin incorporation and westen blot with anti puromycin antibody. ER stress induced by adding 12 uM tunicamycin to tissue incubation solution. B. Quantification of ratio of in-lane puromycin signal/ponceau-labeled signal. n=8 from 4 biological replicates. * p<0.03 ** p<0.008 by 2-tailed Mann-Whitney rank sum test. C. *CenG1a^Δ9^* homozygous adult flies have reduced lifespan. Kaplan-Meier survival curves with 95% confidence interval shaded between dashed lines. Numbers: *w^1118^* 149 flies, 15 vials. *CenG1a^Δ9^* 184 flies, 15 vials. p< 2×10^-16^ log-rank test. D. *CenG1a^Δ9^* flies are more sensitive to tunicamycin lethality. Significantly fewer *CenG1a^Δ9^* flies survived between 90-114 hours from the start of the treatment. No lethality in DMSO control vials was observed during this time. Numbers: *w^1118^* 146 flies, 13 vials. *CenG1a^Δ9^* 139 flies, 13 vials. p< 5×10^-16^ log-rank test.

We next investigated whether impaired eIF2α-P creates functional impairments to stress responses in *CenG1a* mutants. Longevity studies reveal that *CenG1a* mutants die earlier than their genetic controls under normal rearing conditions (**Fig. 6C**). We sought to understand whether mutant animals’ inability to respond to a subsequent cytotoxic stressor would have important consequences at the level of the whole organism. We found that *CenG1a* mutants are more sensitive to ER stress-induced lethality, with tunicamycin-treated animals dying much earlier than genetic controls (**Fig. 6D**). Our findings illustrate a two-hit scenario wherein variants in *CenG1a* increases animals’ susceptibility to a subsequent insult.

## Discussion

We found evidence for an expanded endolysosomal compartment in CenG1a mutant. Accordingly, we also identified increased autophagy and eIF2α-P under basal conditions, suggesting chronic activation of the cytotoxic stress response. Autophagic flux was preserved in mutants, suggesting that lysosomal function was unaffected, but mutant animals had a decreased capacity to respond to “second hit” cytotoxic stressors. When challenged with such a stressor, mutants exhibited diminished survival.

Heterozygous deletions and predicted damaging missense variants in AGAP1 are associated with a mixed neurodevelopmental phenotype that includes autism spectrum disorder and cerebral palsy. *AGAP1* patient phenotypes overlap with those caused by variants in AP3-complex member *AP3D1. AP3D1* variants cause Hermansky-Pudlak syndrome-10. The patient with HP-10 shares several features with *AGAP1* patients, including global developmental delay, epilepsy, dystonia, hypotonia, poor feeding, diminished cortical volumes and poor myelination detected by neuroimaging, and microcephaly (Ammann et al., 2016). Consistent with 1 *AGAP1* patients reported elsewhere (Jin et al., 2020), the *AP3D1* patient had immunodeficiency and died of septic pneumonia at age 3.5 years. Disruptions to endosomal trafficking, such as the retromer complex, are increasingly being linked to developmental disease including hereditary spastic paraplegia and Ritscher-Schinzel Syndrome, with features spanning neurological, skeletal, and immune systems (Yarwood R, 2020, Saitoh, 2022).

We found decreased synaptic size in *Drosophila* with homozygous null variants in *AGAP1/CenG1a* which was not apparent in a previous study decreasing AGAP1 expression using RNAi knockdown or P-element gene disruption (Homma et al., 2014). It is therefore possible that the increased synaptic release found by Homma et al. may represent a mechanism to compensate for *CenG1a* partial loss of function to maintain synaptic function. We did not test whether AGAP1-mediated Arf1 activity or actin cytoskeleton regulation could underly the axon terminal size phenotype. Thus, the molecular mechanism by which *AGAP1* leads to decreased axon terminal size needs further investigation. It is unlikely that decreased axon terminal size or increased synaptic release is due to increased autophagy; increased autophagy is associated with increased synaptogenesis in *Drosophila* and decreased synaptic release (Shen et al., 2015). Therefore, we conclude the autophagy increase we observe at baseline is likely to be a compensatory mechanism.

In addition to an increased rate of basal autophagy, we observed an increase in basal eIF2α-P. Both indicate an activation of stress-response pathways in the absence of external provocation. This argues that this basal activation is instead triggered in response to endogenous factors – the genetic variant of *AGAP1/CenG1a* and the dysregulation of endolysosomal trafficking that ensues.

Although the activation of these elements of the integrated stress response likely serves as a compensatory mechanism, this compensation is only partial, as *CenG1a^Δ9^* mutants show diminished lifespans. Furthermore, this leaves *CenG1a* mutants with a diminished capacity to respond to additional cytotoxic stressors, as mutant animals fail to respond further to a subsequent provocation (amino acid deprivation or tunicamycin). This reduced ability to further activate autophagy or regulate protein synthesis when challenged by exogenous insults is associated with increased mortality. Consistent with the idea of variants in *AGAP1* increasing sensitivity to stress, we noted one of the patients in this cohort had a clear history of an adverse advent that may have also contributed to their disorder. We further identified 4/6 previously published cerebral palsy patients with *AGAP1* variants also had an adverse event in their medical history (Chopra et al., 2022, Jin et al., 2020, van Eyk et al., 2019). This further suggests this gene may contribute to disease in a gene-environment interaction by increasing sensitivity to diverse stressors in some cases. Our findings indicate the gene-environment interaction is not limited to perturbations of protein trafficking as we showed that starvation and ER stress both elicit phenotypes. The integrated stress response may thus represent a point of convergence that connects several forms of genetic and environmental insults. Although this will require additional experimental validation, we suspect that this phenomenon is not specific to *AGAP1/CenG1a* and in fact may be broadly applicable to a variety of gene-environment interactions that contribute to human neurodevelopmental disorders.

## Materials and Methods

### Patient recruitment and data collection

Patients with variants in *AGAP1* were identified through GeneMatcher (Sobreira et al., 2015). Patients were sequenced as part of their medical care. The 2q37.2 deletion limited to *AGAP1* was detected by microarray. Microdeletions were identified by array-CGH (Human Sureprint 2×105K, Agilent technologies) and confirmed with semiquantitative PCR. The missense variant was identified by trio-based whole exome sequencing on DNA derived from blood as previously described (Bayat et al., 2022). Written informed consents for sharing phenotypes were obtained through the treating clinician.

### Drosophila genetics and rearing

*Drosophila melanogaster* were reared on a standard cornmeal, yeast, sucrose food from the BIO5 media facility, University of Arizona. Stocks for experiments were reared at 25°C, 60-80% relative humidity with 12:12 light/dark cycle. Cultures for controls and mutants were maintained with the same growth conditions, especially the density of animals within the vial. A mix of males and females were used for all experiments.

*CenG1a^Δ9^* is a null allele generated by ends-out gene targeting (Gündner et al., 2014) and was a kind gift from M. Hoch. *Df(2L)BSC252*, *w^1118^* and Canton(S) were acquired from Bloomington Stock center (NIH P0OD018537). *CenG1a^Δ9^* was backcrossed with *w^1118^* for 3 generations and w^1118^ was outcrossed with Canton-S and backcrossed with w^1118^ for 12 generations while selecting for red eyes.

### Immunohistochemistry and imaging at the neuromuscular junction

Imaging of neuromuscular junction of 3^rd^ instar wandering larva performed as previously described (Estes et al., 2011). Briefly, a larval filet was dissected in HL-3 Ca++-free saline before fixation in Ca++ free 4% PFA. Washes were performed with PBS with or without 0.1% Triton-X. Blocking before and during primary and secondary antibody steps used PBST with 5% normal goat serum, 2% BSA. Primary antibodies (**Supplemental Table 1**) were incubated overnight at 4°C including DSHB hybridoma monocolonal antibodies: 1:10 mouse α Rab7, 1:200 rabbit α Arl8, 1:400 mouse α DLG (4F3). DSHB antibodies deposited by Munro, S (Rab7, Arl8) and Goodman, C (4F3). Secondary antibodies were incubated for 1.5 hours at RT and included: 1:400 goat α mouse Cy3, goat α rabbit Cy3 or donkey α rabbit Alexa488 (ThermoFisher). Neuronal membranes were visualized with 1:100 goat α HRP-Alexa 647 (Jackson ImmunoResearch) added with secondary antibody; HRP=horseradish peroxidase. 1:300 phallodin-488 in PBS (Molecular Probes) was added as the final wash before mounting.

Imaging was performed with a 710 Zeiss confocal using 1.0 μm Z-stack at 63x of the 1b 6/7 muscle in the A3 abdominal segment. Image parameters (laser gain and intensity, resolution, zoom) were kept constant between images of the same session; alternating between control and mutants. Maximum intensity projections were created for a given field of view (FOV) using all parts of the Z-stack that included the axon terminal (defined by HRP staining) for all channels. Projections converted to greyscale from RGB in Photoshop without adjustments to pixel intensity. Non 6/7 1b HRP staining was identified based on shape/size/HRP staining intensity and manually removed.

Images were analyzed using ImageJ software v. 1.50i (National Institutes of Health). Images from individual channels were converted to black and white using the threshold feature with the threshold held uniform for images within an imaging session. Rab and Arl area was measured using Analyze Particles with a mask created from the HRP image to define and measure the neuron-only area. NMJ area was measured by drawing around the area encompassed by both muscles and neuron in the FOV. Pearson colocalization coefficient was determined using coloc2 plugin from Fiji using the HRP channel to provide a mask for Rab7 and Arl8 channels. Orthogonal projections were converted to 8-bit black and white tiffs without thresholding. Boutons were counted manually from 63x images stitched over the entire neuron area. Statistics and graphs were created using R v. 4.1.0. For boxplots, boxes represent 25^th^ and 75^th^ percentiles; whiskers represent 10^th^ and 90^th^ percentiles.

### Western blotting and quantification

Protein samples were prepared from 10-20 males and females in protein extraction buffer plus 1% protease inhibitor (ThermoFisher) and 1% phosphatase inhibitor (Sigma) as previously described (Emery, 2007). Western blotting was performed according to standard methods with detection on a 0.2 μm nitrocellulose membrane; antibodies detailed in **Supplemental Table 1**. For eIF2α phosphorylation antibody was imaged first, then the membrane was washed and reprobed for total eIF2α. Antibodies used for autophagy: 1:500 rabbit α Ref(2)P (Abcam 178440) normalized using Ponceau staining (Santa Cruz Biotechnology sc-301558) or 1:2000 mouse α beta actin (Abcam 8224) and 1:2000 rabbit α Atg8 (Sigma ABC974). Antibodies used for eIF2α-P: 1:1000 rabbit α phospho-S51 (Cell Signaling 3597), 1:500 rabbit α eIF2S1 (Abcam 4837). Newly synthesized proteins labeled with 1:1000 mouse α puromycin (Kerafast, EQ0001) and normalized to Ponceau staining. Secondary antibodies: 1:10000 goat α rabbit or goat α mouse ECL (GE healthcare NA931, NA934).

Chemoluminescence was captured using a FluorChem Imager (Biotechne) and quantified with Image studio Lite (LiCor). Ref(2), LC3II/LC3-I ratio, and eIF2α phosphorylated/total ratio was normalized to wild-type or no treatment control for each biological replicate. The same size area was used to measure total protein and puromycin signal for each lane to reduce variability. At least 2 technical replicates of each sample were used for quantification. Differences between genotypes were calculated using 2 tailed paired t-test statistic and graphs were generated using R v. 4.1.0. Detailed statistics in **Supplemental Table 2**.

### Drosophila stress paradigms and drug treatment

For tunicamycin exposure, *w^1118^* and *CenG1A^Δ9^* male and female adults at 3 days post-eclosion were anesthetized on ice and pooled into vials filled with 1.3% agar, 1% sucrose with 12 μM tunicamycin (Sigma), with a subset retained for immediate protein extraction. Flies were anesthetized on ice at 4 and 24 hours for protein extraction.

Larva were collected as second instar larva and washed in water. For larval starvation, >20 male and female larva were added to petri dishes made with 1.3% agar and 1% sucrose with or without baker’s yeast supplement and returned to 25°C incubator. Larva were collected after 24 hours; immobile or pupated animals were excluded. Chloroquine treated food was prepared as described by Zirin et al., 2013 at a concentration of 3 mg/mL (Sigma). Larva were collected as above and placed on food mixed with either chloroquine or water as a control for 24 hours and then collected for protein extraction. Successful uptake of drug was confirmed by examining larval gut for bromophenol blue that was mixed into food.

Puromycin labeling performed as described by Deliu et al., 2017. Briefly, 3^rd^ instar larva were floated out of food with 20% sucrose, washed in water, and inverted in batches of 15-20 mixed male and female. Larva were incubated in Schneider’s insect medium (Thermo) with either 12 uM tunicamycin or DMSO (Sigma) control for 3 hours at 25°C. 5 μg/mL of puromycin (Fisher) was added and incubated at 25°C for 40-60 minutes before protein extraction.

### Adult survival

*Drosophila* were collected as late-stage pupae, separated by sex, and sequestered in standard food vials with ~20 animals/vial. The number of alive animals were monitored daily, with animals transferred to fresh food once per week. For tunicamycin lethality, adults were sequestered in vials with 1.3% agar, 1% sucrose with 12 μM tunicamycin or DMSO as a control and monitored twice daily (Chow et al., 2013). Survival library in R v. 4.1.0 was used to generate Kaplan-Meier plots and the log-rank test statistic.

## Acknowledgements

We appreciate the participation by patients and their families for these studies. We acknowledge F. Nowlen and J. Liu for assistance with fly rearing and survival monitoring and A. Musmacker for western blotting. Stocks obtained from the Bloomington Drosophila Stock Center (NIH P40OD018537) were used in this study. The monoclonal antibodies, developed by S. Munro and C. Goodman were obtained from the Developmental Studies Hybridoma Bank, created by the NICHD of the NIH and maintained at The University of Iowa, Department of Biology, Iowa City, IA 52242. Biomedical Imaging Core microscopy facilities at the University of Arizona College of Medicine – Phoenix were used in these studies.

## Competing interests

No competing interests declared.

## Funding

These studies were supported by National Institutes of Health R01NS106298 awarded to MCK.

## Data availability statement

The datasets generated during and/or analyzed during the current study are available from the corresponding author on reasonable request.

## Notes

### Competing Interest Statement

The authors have declared no competing interest.

## References

Ahluwalia, J. K., Hariharan, M., Bargaje, R., Pillai, B. & Brahmachari, V. 2009. Incomplete penetrance and variable expressivity: is there a microRNA connection? Bioessays, 31, 981–92.

Ammann, S., Schulz, A., Krägeloh-Mann, I., Dieckmann, N., Niethammer, K., Fuchs, S., Eckl, K., Plank, R., Werner, R., Altmüller, J., Thiele, H., Nürnberg, P., Bank, J., Strauss, A, Von Bernuth, H., Zur Stadt, U., Grieve, S., Griffiths, G., Lehmberg, K., Hennies, H. & Ehl, S. 2016. Mutations in AP3D1 associated with immunodeficiency and seizures define a new type of Hermansky-Pudlak syndrome. Blood, 127, 997–1006.

Amodio, G., Moltedo, O., Fasano, D., Zerillo, L., Oliveti, M., Di Pietro, P., Faraonio, R., Barone, P., Pellecchia, M., De Rosa, A., De Michele, G., Polishchuk, E., Polishchuk, R., Bonifati, V., Nitsch, L., Pierantoni, G., Renna, M., Criscuolo, C., Paladino, S. & Remondelli, P. 2019. PERK-Mediated Unfolded Protein Response Activation and Oxidative Stress in PARK20 Fibroblasts. Front Neurosci, 13, 673.

Arnold, M., Cross, R., Singleton, K., Zlatic, S., Chapleau, C., Mullin, A., Rolle, I., Moore, C., Theibert, A., Pozzo-Miller, L., Faundez, V. & Larimore, J. 2016. The Endosome Localized Arf-GAP AGAP1 Modulates Dendritic Spine Morphology Downstream of the Neurodevelopmental Disorder Factor Dysbindin. Front Cell Neurosci, 10.

B’Chir, W., Maurin, A., Carraro, V., Averous, J., Jousse, C., Muranishi, Y., Parry, L., Stepien, G., Fafournoux, P. & Bruhat, A. 2013. The eIF2α/ATF4 pathway is essential for stress-induced autophagy gene expression. Nucleic Acids Res, 41, 7683–99.

Bayat, A., Fenger, C. D., Techlo, T. R., Højte, A. F., Nørgaard, I., Hansen, T. F., Rubboli, G., Møller, R. S. & Group, D. 2022. Impact of Genetic Testing on Therapeutic Decision-Making in Childhood-Onset Epilepsies-a Study in a Tertiary Epilepsy Center. Neurotherapeutics, 19, 1353–1367.

Bendor, J., Lizardi-Ortiz, J., Westphalen, R., Brandstetter, M., Hemmings, H. J., Sulzer, D., Flajolet, M. & Greengard, P. 2010. AGAP1/AP-3-dependent endocytic recycling of M5 muscarinic receptors promotes dopamine release. EMBO J, 29, 2813–26.

Brunet, T., Jech, R., Brugger, M., Kovacs, R., Alhaddad, B., Leszinski, G., Riedhammer, K. M., Westphal, D. S., Mahle, I., Mayerhanser, K., Skorvanek, M., Weber, S., Graf, E., Berutti, R., Necpál, J., Havránková, P., Pavelekova, P., Hempel, M., Kotzaeridou, U., Hoffmann, G. F., Leiz, S., Makowski, C., Roser, T., Schroeder, S. A., Steinfeld, R., Strobl-Wildemann, G., Hoefele, J., Borggraefe, I., Distelmaier, F., Strom, T. M., Winkelmann, J., Meitinger, T., Zech, M. & Wagner, M. 2021. De novo variants in neurodevelopmental disorders-experiences from a tertiary care center. Clin Genet, 100, 14–28.

Chapuy, B., Tikkanen, R., Mühlhausen, C., Wenzel, D., Von Figura, K. – Höning, S. 2008. AP-1 and AP-3 mediate sorting of melanosomal and lysosomal membrane proteins into distinct post-Golgi trafficking pathways. Traffic, 9, 1157–72.

Chopra, M., Gable, D. L., Love-Nichols, J., Tsao, A., Rockowitz, S., Sliz, P., Barkoudah, E., Bastianelli, L., Coulter, D., Davidson, E., Degusmao, C., Fogelman, D., Huth, K., Marshall, P., Nimec, D., Sanders, J. S., Shore, B. J., Snyder, B., Stone, S. S. D., Ubeda, A., Watkins, C., Berde, C., Bolton, J., Brownstein, C., Costigan, M., Ebrahimi-Fakhari, D., Lai, A., O’Donnell-Luria, A., Paciorkowski, A. R., Pinto, A., Pugh, J., Rodan, L., Roe, E., Swanson, L., Zhang, B., Kruer, M. C., Sahin, M., Poduri, A. & Srivastava, S. 2022. Mendelian etiologies identified with whole exome sequencing in cerebral palsy. Ann Clin Transl Neurol, 9, 193–205.

Chow, C., Wolfner, M. & Clark, A. 2013. Using natural variation in Drosophila to discover previously unknown endoplasmic reticulum stress genes. Proc Natl Acad Sci U S A, 110, 9013–8.

Cukier, H., Dueker, N., Slifer, S., Lee, J., Whitehead, P., Lalanne, E., Leyva, N., Konidari, I., Gentry, R., Hulme, W., Booven, D., Mayo, V., Hofmann, N., Schmidt, M., Martin, E., Haines, J., Cuccaro, M., Gilbert, J. & Pericak-Vance, M. 2014. Exome sequencing of extended families with autism reveals genes shared across neurodevelopmental and neuropsychiatric disorders. Mol Autism, 5, 1.

Cukierman, E., Huber, I., Rotman, M. & Cassel, D. 1995. The ARF1 GTPase-activating protein: zinc finger motif and Golgi complex localization. Science, 270, 1999–2002.

Davidson, A., Humphreys, D., Brooks, A., Hume, P. & Koronakis, V. 2015. The Arf GTPase-activating protein family is exploited by Salmonella enterica serovar Typhimurium to invade nonphagocytic host cells. mBio, 6, :e02253–14.

Emery, P. 2007. Protein extraction from Drosophila heads. Methods Mol Biol, 362, 375–7.

Estes, P., Boehringer, A., Zwick, R., Tang, J., Grigsby, B. & Zarnescu, D. 2011. Wild-type and A315T mutant TDP-43 exert differential neurotoxicity in a Drosophila model of ALS. Hum Mol Genet, 20, 2308–2321.

Fahey, M. C., Maclennan, A. H., Kretzschmar, D., Gecz, J. & Kruer, M. C. 2017. The genetic basis of cerebral palsy. Dev Med Child Neurol, 59, 462–469.

Geschwind, D. H. 2011. Genetics of autism spectrum disorders. Trends Cogn Sci, 15, 409–16.

Gündner, A., Hahn, I., Sendscheid, O., Aberle, H. & Hoch, M. 2014. The PIKE homolog Centaurin gamma regulates developmental timing in Drosophila. PLoS One, 9, e97332.

Heuvingh, J., Franco, M., Chavrier, P. & Sykes, C. 2007. ARF1-mediated actin polymerization produces movement of artificial vesicles. Proc Natl Acad Sci U S A, 104, 16928–33.

Homma, M., Nagashima, S., Fukuda, T., Yanagi, S., Miyakawa, H., Suzuki, E. & Morimoto, T. 2014. Downregulation of Centaurin gamma1A increases synaptic transmission at Drosophila larval neuromuscular junctions. Eur J Neurosci, 40, 3158–70.

Humeau, J., Leduc, M., Cerrato, G., Loos, F., Kepp, O. & Kroemer, G. 2020. Phosphorylation of eukaryotic initiation factor-2α (eIF2α) in autophagy. Cell Death Dis, 11, 433.

Jacomin, A. C., Fauvarque, M. O. & Taillebourg, E. 2016. A functional endosomal pathway is necessary for lysosome biogenesis in Drosophila. BMC Cell Biol, 17, 36.

Jin, S. C., Lewis, S. A., Bakhtiari, S., Zeng, X., Sierant, M. C., Shetty, S., Nordlie, S. M., Elie, A., Corbett, M. A., Norton, B. Y., Van Eyk, C. L., Haider, S., Guida, B. S., Magee, H., Liu, J., Pastore, S., Vincent, J. B., Brunstrom-Hernandez, J., Papavasileiou, A., Fahey, M. C., Berry, J. G., Harper, K., Zhou, C., Zhang, J., Li, B., Zhao, H., Heim, J., Webber, D. L., Frank, M. S. B., Xia, L., Xu, Y., Zhu, D., Zhang, B., Sheth, A. H., Knight, J. R., Castaldi, C., Tikhonova, I. R., LÓPez-Giráldez, F., Keren, B., Whalen, S., Buratti, J., Doummar, D., Cho, M., Retterer, K., Millan, F., Wang, Y., Waugh, J. L., Rodan, L., Cohen, J. S., Fatemi, A., Lin, A. E., Phillips, J. P., Feyma, T., Maclennan, S. C., Vaughan, S., Crompton, K. E., Reid, S. M., Reddihough, D. S., Shang, Q., Gao, C., Novak, I., Badawi, N., Wilson, Y. A., Mcintyre, S. J., Mane, S. M., Wang, X., Amor, D. J., Zarnescu, D. C., Lu, Q., Xing, Q., Zhu, C., Bilguvar, K., Padilla-Lopez, S., Lifton, R. P., Gecz, J., Maclennan, A. H. & Kruer, M. C. 2020. Mutations disrupting neuritogenesis genes confer risk for cerebral palsy. Nat Genet, 52, 1046–1056.

Klionsky, D. J., Abeliovich, H., Agostinis, P., Agrawal, D. K., Aliev, G., Askew, D. S., Baba, M., Baehrecke, E. H., Bahr, B. A., Ballabio, A., Bamber, B. A., Bassham, D. C., Bergamini, E., Bi, X., Biard-Piechaczyk, M., Blum, J. S., Bredesen, D. E., Brodsky, J. L., Brumell, J. H., Brunk, U. T., Bursch, W., Camougrand, N., Cebollero, E., Cecconi, F., Chen, Y., Chin, L. S., Choi, A., Chu, C. T., Chung, J., Clarke, P. G., Clark, R. S., Clarke, S. G., ClavÉ, C., Cleveland, J. L., Codogno, P., Colombo, M. I., Coto-Montes, A., Cregg, J. M., Cuervo, A. M., Debnath, J., Demarchi, F., Dennis, P. B., Dennis, P. A., Deretic, V., Devenish, R. J., Di Sano, F., Dice, J. F., Difiglia, M., Dinesh-Kumar, S., Distelhorst, C. W., Djavaheri-Mergny, M., Dorsey, F. C., Dröge, W., Dron, M., Dunn, W. A., JR., Duszenko, M., Eissa, N. T., Elazar, Z., Esclatine, A., Eskelinen, E. L., FÉSüs, L., Finley, K. D., Fuentes, J. M., Fueyo, J., Fujisaki, K., Galliot, B., Gao, F. B., Gewirtz, D. A., Gibson, S. B., Gohla, A., Goldberg, A. L., Gonzalez, R., González-EstÉvez, C., Gorski, S., Gottlieb, R. A., Häussinger, D., He, Y. W., Heidenreich, K., Hill, J. A., Høyer-Hansen, M., Hu, X., Huang, W. P., Iwasaki, A., Jäättelä, M., Jackson, W. T., Jiang, X., Jin, S., Johansen, T., Jung, J. U., Kadowaki, M., Kang, C., Kelekar, A., Kessel, D. H., Kiel, J. A., Kim, H. P., Kimchi, A., Kinsella, T. J., Kiselyov, K., Kitamoto, K., Knecht, E., et al. 2008. Guidelines for the use and interpretation of assays for monitoring autophagy in higher eukaryotes. Autophagy, 4, 151–75.

Leroy, C., Landais, E., Briault, S., David, A., Tassy, O., Gruchy, N., Delobel, B., GrÉgoire, M., Leheup, B., Taine, L., Lacombe, D., Delrue, M., Toutain, A., Paubel, A., Mugneret, F., Thauvin-Robinet, C., Arpin, S., Le Caignec, C., Jonveaux, P., Beri, M., Leporrier, N., Motte, J., Fiquet, C., Brichet, O., Mozelle-Nivoix, M., Sabouraud, P., Golovkine, N., Bednarek, N., Gaillard, D. & Doco-Fenzy, M. 2013. The 2q37-deletion syndrome: an update of the clinical spectrum including overweight, brachydactyly and behavioural features in 14 new patients. Eur J Hum Genet, 21, 602–12.

Lincoln, B. L., 2ND, Alabsi, S. H., Frendo, N., Freund, R. & Keller, L. C. 2015. Drosophila Neuronal Injury Follows a Temporal Sequence of Cellular Events Leading to Degeneration at the Neuromuscular Junction. J Exp Neurosci, 9, 1–9.

Madsen, A. M., Hodge, S. E. & Ottman, R. 2011. Causal models for investigating complex disease: I. A primer. Hum Hered, 72, 54–62.

Mahjani, B., De Rubeis, S., Gustavsson Mahjani, C., Mulhern, M., Xu, X., Klei, L., Satterstrom, F. K., Fu, J., Talkowski, M. E., Reichenberg, A., Sandin, S., Hultman, C. M., Grice, D. E., Roeder, K., Devlin, B. & Buxbaum, J. D. 2021. Prevalence and phenotypic impact of rare potentially damaging variants in autism spectrum disorder. Mol Autism, 12, 65.

Moreno-De-Luca, A., Millan, F., Pesacreta, D. R., Elloumi, H. Z., Oetjens, M. T., Teigen, C., Wain, K. E., Scuffins, J., Myers, S. M., Torene, R. I., Gainullin, V. G., Arvai, K., Kirchner, H. L., Ledbetter, D. H., Retterer, K. & Martin, C. L. 2021. Molecular Diagnostic Yield of Exome Sequencing in Patients With Cerebral Palsy. Jama, 325, 467–475.

Nie, Z., Boehm, M., Boja, E., Vass, W., Bonifacino, J., Fales, H. & Randazzo, P. 2003. Specific regulation of the adaptor protein complex AP-3 by the Arf GAP AGAP1. Dev Cell, 5, 513–21.

Pacault, M., Nizon, M., Pichon, O., Vincent, M., Le Caignec, C. & Isidor, B. 2018. A de novo 2q37.2 deletion encompassing AGAP1 and SH3BP4 in a patient with autism and intellectual disability. Eur J Med Genet., 62, 103586.

Pakos-Zebrucka, K., Koryga, I., Mnich, K., Ljujic, M., Samali, A. & Gorman, A. 2016. The integrated stress response. EMBO Rep, 17, 1374–1395.

Pircs, K., Nagy, P., Varga, A., Venkei, Z., Erdi, B., Hegedus, K. & Juhasz, G. 2012. Advantages and limitations of different p62-based assays for estimating autophagic activity in Drosophila. PLoS One, 7, e44214.

Saila, S., Gyanwali, G. C., Hussain, M., Gianfelice, A. & Ireton, K. 2020. The Host GTPase Arf1 and Its Effectors AP1 and PICK1 Stimulate Actin Polymerization and Exocytosis To Promote Entry of Listeria monocytogenes. Infect Immun, 88.

Saitoh, S. 2022. Endosomal Recycling Defects and Neurodevelopmental Disorders. Cells, 11.

Shearer, L. & Petersen, N. 2019. Distribution and Co-localization of endosome markers in cells. Heliyon, 5, e02375.

Shen, D. N., Zhang, L. H., Wei, E. Q. & Yang, Y. 2015. Autophagy in synaptic development, function, and pathology. Neurosci Bull, 31, 416–26.

Sobreira, N., Schiettecatte, F., Valle, D. & Hamosh, A. 2015. GeneMatcher: A Matching Tool for Connecting Investigators with an Interest in the Same Gene. Hum Mutat, 36, 928–30.

Ugur B, C. K., Bellen HJ 2016. Drosophila tools and assays for the study of human disease. Dis Model Mech, 9, 235–44.

Van Eyk Cl, C. M., Frank MSB, Webber DL, Newman, M, Berry JG, Harper K, Haines BP, Mcmichael G, Woenig JA, Maclennan AH, Gecz J 2019. Targeted resequencing identifies genes with recurrent variation in cerebral palsy. NPJ Genom Med, 4, 27.

Van Eyk, C. L., Webber, D. L., Minoche, A. E., PÉRez-Jurado, L. A., Corbett, M. A., Gardner, A. E., Berry, J. G., Harper, K., Maclennan, A. H. & Gecz, J. 2021. Yield of clinically reportable genetic variants in unselected cerebral palsy by whole genome sequencing. Npj Genom Med, 6, 74.

Vasin, A., Zueva, L., Torrez, C., Volfson, D., Littleton, J. T. & Bykhovskaia, M. 2014. Synapsin regulates activity-dependent outgrowth of synaptic boutons at the Drosophila neuromuscular junction. J Neurosci, 34, 10554–63.

Xiong R, S. D., Ross D 2013. The activation sequence of cellular protein handling systems after proteasomal inhibition in dopaminergic cells. Chem Biol Interact, 204, 116–24.

Yarwood R, H. J., Woodman PG, Lowe M 2020. Membrane trafficking in health and disease. Dis Model Mech, 13, dmm043448.

Yuen, C. R. K., Merico, D., Bookman, M., J, L. H., Thiruvahindrapuram, B., Patel, R. V., Whitney, J., Deflaux, N., Bingham, J., Wang, Z., Pellecchia, G., Buchanan, J. A., Walker, S., Marshall, C. R., Uddin, M., Zarrei, M., Deneault, E., D’Abate, L., Chan, A. J., Koyanagi, S., Paton, T., Pereira, S. L., Hoang, N., Engchuan, W., Higginbotham, E. J., Ho, K., Lamoureux, S., Li, W., Macdonald, J. R., Nalpathamkalam, T., Sung, W. W., Tsoi, F. J., Wei, J., Xu, L., Tasse, A. M., Kirby, E., Van Etten, W., Twigger, S., Roberts, W., Drmic, I., Jilderda, S., Modi, B. M., Kellam, B., Szego, M., Cytrynbaum, C., Weksberg, R., Zwaigenbaum, L., Woodbury-Smith, M., Brian, J., Senman, L., Iaboni, A., Doyle-Thomas, K., Thompson, A., Chrysler, C., Leef, J., Savion-Lemieux, T., Smith, I. M., Liu, X., Nicolson, R., Seifer, V., Fedele, A., Cook, E. H., Dager, S., Estes, A., Gallagher, L., Malow, B. A., Parr, J. R., Spence, S. J., Vorstman, J., Frey, B. J., Robinson, J. T., Strug, L. J., Fernandez, B. A., Elsabbagh, M., Carter, M. T., Hallmayer, J., Knoppers, B. M., Anagnostou, E., Szatmari, P., Ring, R. H., Glazer, D., Pletcher, M. T. & Scherer, S. W. 2017. Whole genome sequencing resource identifies 18 new candidate genes for autism spectrum disorder. Nat Neurosci, 20, 602–611.

Zirin, J., Nieuwenhuis, J. & Perrimon, N. 2013. Role of autophagy in glycogen breakdown and its relevance to chloroquine myopathy. PLoS Biol, 11, e1001708.

