## Supplementary material for "AGAP1-associated endolysosomal trafficking abnormalities link gene-environment interactions in a neurodevelopmental disorder": Table S1

| Antibody/stain | Species | Concentration | Application | Company | Lot(s) # |
| --- | --- | --- | --- | --- | --- |
| Rab7 | M | 1:10 | NMJ IHC | DSHB | 9-26-16 |
| Arl8 polyclonal supernatent | R | 1:200 | NMJ IHC | DSHB | 6/9/16 |
| DLG (4F3) | M | 1:400 | NMJ IHC | DSHB | 2/18/16 |
| Anti-mouse Cy3 | G | 1:400 | NMJ IHC | ThermoFisher | A10521 1425618 |
| Anti-rabbit Cy3 | G | 1:400 | NMJ IHC | ThermoFisher | A10520  1633861 |
| Anti-rabbit 488 | D | 1:400 | NMJ IHC | ThermoFisher | A11008  1515529 |
| Anti-HRP 647 | G | 1:100 | NMJ IHC | Jackson ImmunoResearch | 126323 |
| Phalloidin 488 | - | 1:300 | NMJ IHC | Molecular Probes | 1903540  2160010 |
| Ref(2)P | R | 1:500 | WB | Abcam 178440 | GR3389640 |
| Beta actin | M | 1:2000 | WB | Abcam 8224 | GR14272-3 |
| Beta tubulin | R | 1:2000 | WB | Abcam 6046 | GR3376491-1 |
| Atg8a | R | 1:2000 | WB | Sigma ABC974 | 3308314 |
| Phosphor-S51 eIF2S | R | 1:1000 | WB | Cell Signaling 3597 | 12 |
| eIF2S1 | R | 1:500 | WB | Abcam 26197 | GR3183673-1  GR3325905 |
| Puromycin monoclonal | M | 1:1000 | WB | Kerafast EQ0001 | 3RH11 |
| Rabbit ECL | G | 1:10000 | WB | GE healthcare NA931 | 17473046 |
| Mouse ECL | G | 1:10000 | WB | GE Healthcare NA934 | 17170583 |

Supplemental Table 1: Antibodies and biological reagents used in studies

Additional biologicals:

Normal goat serum Abcam ab7481 lot GR3188160

Schneider’s insect media (Thermo) 21720024 lot 1894706
