## Supplementary material for "AGAP1-associated endolysosomal trafficking abnormalities link gene-environment interactions in a neurodevelopmental disorder": Table S2

|  | Test used/rationale | Genotype comparison | Result (p-value) |
| --- | --- | --- | --- |
| Fig 2B | 2-tailed Mann-Whitney rank sum test. More conservative non-parametric statistic chosen due to unknown if variance is the same between genotypes due to immunostaining and microscopy methods. | W1118 vs Δ9 homozygote | 0.03 |
| Fig 2B |  | W1118 vs Δ9/Df hemizygote | 0.004 |
| Fig 2B |  | Δ9 homozygote vs Δ9/Df hemizygote | n.s. |
| Fig 2C |  | W1118 vs Δ9 homozygote | n.s. |
| Fig 2C |  | W1118 vs Δ9/Df hemizygote | 0.009 |
| Fig 2C |  | Δ9 homozygote vs Δ9/Df hemizygote | 0.001 |
| Fig 2D |  | W1118 vs Δ9 homozygote | n.s. |
| Fig 2D |  | W1118 vs Δ9/Df hemizygote | n.s. |
| Fig 2D |  | Δ9 homozygote vs Δ9/Df hemizygote | n.s. |
| Fig 2E |  | W1118 vs Δ9 homozygote | n.s. |
| Fig 2E |  | W1118 vs Δ9/Df hemizygote | n.s. |
| Fig 2E |  | Δ9 homozygote vs Δ9/Df hemizygote | n.s. |
| Fig 3C | 2-tailed Mann-Whitney rank sum test. More conservative non-parametric statistic chosen due to unknown if variance is the same between genotypes due to immunostaining and microscopy methods. | W1118 vs Δ9 homozygote NMJ | n.s. |
| Fig 3C |  | W1118 vs Δ9 homozygote neuron | 0.02 |
| Fig 3C’ |  | W1118 vs Δ9 homozygote NMJ | 9.596e-05 |
| Fig 3C’ |  | W1118 vs Δ9 homozygote neuron | n.s. |
| Fig 3C’’ |  | W1118 vs Δ9 homozygote NMJ | 0.03 |
| Fig 3C’’ |  | W1118 vs Δ9 homozygote neuron | 0.01 |
| Fig 3F |  | W1118 vs Δ9 homozygote NMJ | 0.003 |
| Fig 3F |  | W1118 vs Δ9 homozygote neuron | 0.02 |
| Fig 3F’ |  | W1118 vs Δ9 homozygote NMJ | n.s. |
| Fig 3F’ |  | W1118 vs Δ9 homozygote neuron | n.s. |
| Fig 3F’’ |  | W1118 vs Δ9 homozygote NMJ | 0.02 |
| Fig 3F’’ |  | W1118 vs Δ9 homozygote neuron | 0.03 |
| Fig 3G |  | W1118 vs Δ9 homozygote | 0.02 |
| Fig 4B | p<0.05 by 2-tailed paired t-test. T-test chosen as biological samples prepared in parallel and normalized to genetic and treatment control. | *w^1118^* control vs starve | n.s. |
| Fig 4B |  | Control *w^1118^* vs *Δ9* homozygote | 0.04 |
| Fig 4B |  | *Δ9* homozygote control vs starve | 0.03 |
| Fig 4B |  | Starve *w^1118^* vs *Δ9* homozygote | n.s. |
| Fig 4C |  | *w^1118^* control vs starve | n.s. |
| Fig 4C |  | Control *w^1118^* vs *Δ9* homozygote | 0.02 |
| Fig 4C |  | *Δ9* homozygote control vs starve | n.s. |
| Fig 4C |  | Starve *w^1118^* vs *Δ9* homozygote | n.s. |
| Fig 4E | p<0.05 by 2-tailed paired t-test. T-test chosen as biological samples prepared in parallel and normalized to genetic and treatment control. | *w^1118^* control vs chloroquine | 0.009 |
| Fig 4E |  | Control *w^1118^* vs *Δ9* homozygote | 0.02 |
| Fig 4E |  | *Δ9* homozygote control vs chloroquine | 0.03 |
| Fig 4E |  | chloroquine *w^1118^* vs *Δ9* homozygote | n.s. |
| Fig 4F |  | *w^1118^* control vs chloroquine | n.s. |
| Fig 4F |  | Control *w^1118^* vs *Δ9* homozygote | 0.05 |
| Fig 4F |  | *Δ9* homozygote control vs chloroquine | n.s. |
| Fig 4F |  | chloroquine *w^1118^* vs *Δ9* homozygote | n.s. |
| Fig 5B | 2-tailed paired t-test. T-test chosen as biological samples prepared in parallel and normalized to genetic and treatment control. | *w^1118^* vs *Δ9* homozygote 3L |  |
| Fig 5B |  | *w^1118^* vs *Δ9* homozygote 1 day | n.s. |
| Fig 5B |  | *w^1118^* vs *Δ9* homozygote 14 day | 0.003 |
| Fig 5B |  | *w^1118^* 1 day vs 14 days | 0.004 |
| Fig 5B |  | *Δ9* homozygote 1 day vs 14 days | n.s. |
| Fig 5C |  | *w^1118^* control vs starvation | 0.005 |
| Fig 5C |  | Control *w^1118^* vs *Δ9* homozygote | 0.06 |
| Fig 5C |  | *Δ9* homozygote control vs starvation | n.s. |
| Fig 5C |  | starvation *w^1118^* vs *Δ9* homozygote | n.s. |
| Fig 5D |  | *w^1118^* 0 v 4 hour tunicamycin | 0.03 |
| Fig 5D |  | *w^1118^* 0 v 24 hour tunicamycin | 0.03 |
| Fig 5D |  | *w^1118^* 4 v 24 hour tunicamycin | n.s. |
| Fig 5D |  | *Δ9* homozygote 0 v 4 hour tunicamycin | n.s. |
| Fig 5D |  | *Δ9* homozygote 0 v 24 hour tunicamycin | 0.01 |
| Fig 5D |  | *Δ9* homozygote 4 v 24 hour tunicamycin | n.s. |
| Fig 5D |  | *w^1118^* v *Δ9* homozygote 0 hour | n.s. |
| Fig 5D |  | *w^1118^* v *Δ9* homozygote 4 hour | n.s. |
| Fig 5D |  | *w^1118^* v *Δ9* homozygote 24 hour | n.s. |
| Fig 6B | 2-tailed Mann-Whitney rank sum test chosen because using non-normalized data. | *w^1118^* control vs tunicamycin | 0.02 |
| Fig 6B |  | *w^1118^* vs *Δ9* homozygote control | 0.008 |
| Fig 6B |  | *Δ9* control vs tunicamycin | n.s. |
| Fig 6B |  | *w^1118^* vs *Δ9* homozygote tunicamycin | n.s. |
| Fig 6C | Kaplan meyer log-rank test. This is the standard hypothesis-based, nonparametric test to compare the survival distributions of two samples and accounts for the right-hand skew. | *w^1118^* vs *Δ9* homozygote | p< 2x10^-16^ |
| Fig 6D |  | *w^1118^* vs *Δ9* homozygote | p< 5x10^-16^ |

Supplemental table 2: Statistical tests and results from studies.
